# The effect of strain level diversity on robust inference of virus-induced mortality

**DOI:** 10.1101/277517

**Authors:** Stephen J. Beckett, Joshua S. Weitz

**Affiliations:** School of Biological Sciences, Georgia Institute of Technology, Atlanta, GA, USA; School of Physics, Georgia Institute of Technology, Atlanta, GA, USA

## Abstract

Infection and lysis of phytoplankton by viruses affects population dynamics and nutrient cycles within oceanic microbial communities. However, estimating the quantitative rates of viral-induced lysis remains challenging in situ. The modified dilution method is the most commonly utilised empirical approach to estimate virus-induced killing rates of phytoplankton. The lysis rate estimates of the modified dilution method are based on models of virus-host interactions involving only a single virus and a single host population. Here, using modelling approaches, we examine the robustness of the modified dilution method in multi-strain, complex communities. We assume that strains differ in their life history traits, including growth rates (of hosts) and lysis rates (by viruses). We show that trait differences affect resulting experimental dynamics such that lysis rates measured using the modified dilution method may be driven by the fastest replicating strains; which are not necessarily the most abundant in situ. We discuss the implications of using the modified dilution method and alternative dilution-based approaches for estimating viral-induced lysis rates in marine microbial communities.

## 1 INTRODUCTION

Viruses and grazers impact marine microbial populations and biogeochemical cycling. Grazing by micrograzers transfers carbon from primary producers to higher trophic levels and modifies rates of remineralisation and sinking of organic matter (Sherr and Sherr, 2002; Calbet and Landry, 2004). Viral induced cellular lysis may release resources that can be utilised by primary producers in the so-called “viral shunt” (Wilhelm and Suttle, 1999; Suttle, 2007; Weitz and Wilhelm, 2012). Quantifying the rates of grazing and viral lysis on marine microbes is important to understand the dynamics of marine ecosystems.

The dilution method is the prevailing technique used to evaluate the impact of grazers on microbial populations (Landry and Hassett, 1982). We term this the ‘classic’ dilution method (CDiM). The CDiM measures the differential rate of recovery across incubation experiments containing different levels of diluted seawater. The modified dilution method (MDiM) (Evans et al., 2003; Kimmance and Brussaard, 2010) is used to quantify the impact of both grazing and viral lysis on microbial populations. Empirical measurements made via the MDiM suggest that both grazing and viral lysis are important mortality drivers of marine microbes; and that both can be the dominant source of mortality (Tsai et al., 2013; Mojica et al., 2015; Pasulka et al., 2015). Yet, the MDiM can yield negative rates of viral lysis (Pasulka et al., 2015) and there are concerns about the ability to detect viral impact e.g. due to previously infected cells, the length of latent period, and the specificty of viral infection (Jacquet et al., 2005; Kimmance et al., 2007; Kimmance and Brussaard, 2010; Pasulka et al., 2015). Irrespective of the system, the core inference approach underlying the MDiM assumes that ecosystem includes a single, dominant host and viral population. This assumption may limit the application of the MDiM to diverse environments (Fredrickson et al., 2011; Menden-Deuer and Rowlett, 2014; Needham et al., 2017; Aylward et al., 2017).

In this mansucript, we evaluate the potential for the modified dilution method to infer viral lysis in complex multi-strain communities. In doing so, we develop nonlinear dynamic models of virus-host interactions taking place in diluted media. In doing so, we also propose an alternative conceptual approach: a virus dilution method (VDiM) which dilutes only viruses, but not microbes. We find that the VDiM may reduce estimation bias compared to the MDiM, particularly given multi-strain scenarios. In doing so, we provide a theoretical framework to connect empirical measurements with the nonlinear feedbacks and interactions we are trying to infer. We hope that improved estimates of viral lysis rates will deepen understanding of the role of viruses in shaping community structure and biogeochemical fluxes (Brussaard et al., 2008; Weitz et al., 2015; Mateus, 2017).

## 2 MATERIALS AND METHODS

### 2.1 Modified dilution method

The modified dilution method utilises two dilution series: the *classical dilution series* which dilutes phytoplankton and grazers (but not viruses) and the *modified dilution series* which dilutes phytoplankton, grazers and viruses.

To determine rates of viral induced lysis using the modified dilution method two parallel dilution series are created (Figure 1a). Two diluents are created using different filter sizes; the classic filter which excludes phytoplankton and grazers (as in the classical dilution method (Landry and Hassett, 1982)), and another filter small enough to exclude viruses, phytoplankton and grazers. Each dilution series contains several incubation experiments which comprise a different proportion *F* of sampled seawater and (1 — *F*) diluent. Each incubation typically lasts a day, upon which the change in phytoplankton abundance is measured allowing the approximate growth rate within an incubation bottle to be calculated. By plotting apparent growth rate vs. dilution level (*F*), it is possible to draw dilution curves for both the classical and modified dilution series (Figure 1c). The expectation is that the slope of the classical dilution curve is equivalent to the grazing rate (Landry and Hassett, 1982; Kimmance and Brussaard, 2010; Beckett and Weitz, 2017), the intercept of the modified dilution curve at *F* = 0 is the phytoplankton growth rate (Kimmance and Brussaard, 2010), whilst the slope of the modified dilution curve is the sum of both grazing and lysis rates (Kimmance and Brussaard, 2010). Hence, the difference in slopes between the classical and modified dilution curves is expected to be the viral lysis rate.

**Figure 1.**
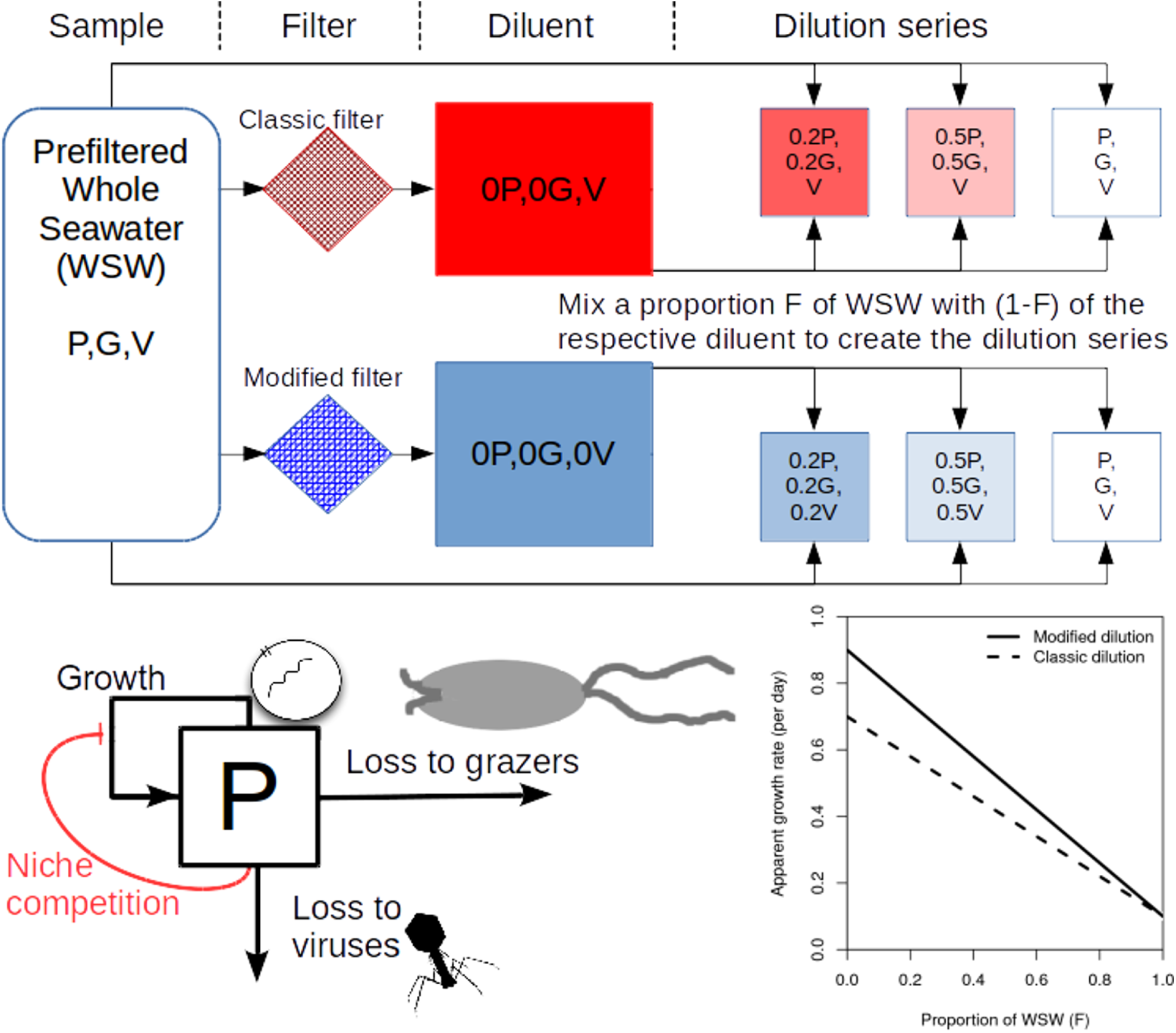
Schematic representing the modified dilution method. (a) scheme to generate the classical dilution series (top), by mixing WSW with a diluent that filters out phytoplankton and grazers; and the modified dilution series (bottom) by mixing WSW with a diluent that filters out viruses, phytoplankton and grazers. (b) A diagram showing growth and mortality processes of phytoplankton P. (c) Idealised apparent growth curves from the classical and modified dilution series. Viral lysis rates are estimated based on the differences between slopes of the classic and modified dilution curves.

### 2.2 Virus dilution method

We propose an alternative conceptual approach to estimating viral lysis. The virus dilution method is comprised of one dilution series. The *virus dilution series* dilutes the ambient levels of viruses (but not phytoplankton or grazers). Each incubation experiment within the viral dilution series contains a proportion *F* of sampled seawater and a proportion (1 — *F*) of seawater from which free viruses are removed. By plotting the apparent growth rates of phytoplankton against the dilution level (*F*) a viral dilution curve can be drawn. Because only the concentration of viruses changes across the virus dilution series, that the slope of the viral dilution curve represents an alternative approach to infer viral lysis rates. We recognise that practical implementation of this method may be challenging and we return to this in the Discussion. However, we can directly apply and assess the viral dilution methodology *in silico* as a means to test the potential accuracy and associated biases of the approach.

### 2.3 In silico dilution experiments

*In silico* dilution experiments were used to estimate viral lysis and compare it against the model input viral lysis rate. Ten dilution levels were used (*F* = 0.1, 0.2, 0.3, 0.4, 0.5, 0.6, 0.7, 0.8, 0.9, 1.0) in each simulation series. Apparent growth rates, 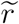, are calculated as:

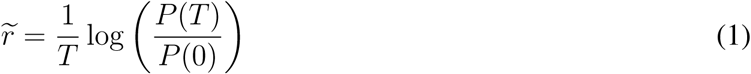

where *T* is the length of incubation and *P*(0) and *P*(*T*) are countable phytoplankton cell densities at the beginning (time 0) and end of the incubation period (time *T*) respectively. Slopes and intercepts of the respective dilution curves are estimated using linear regression. Grazing rate is estimated as the slope of the classical dilution curve and the viral lysis rate is estimated by (a) the difference in slopes between the modified and classical dilution curves and (b) the slope of the viral dilution curve. Bias is calculated by dividing the estimated viral lysis rate by the model input viral lysis rate. Ambient concentrations of phytoplankton and viruses are assumed to be at steady-state. Population dynamics are simulated using package deSolve v1.20 (Soetaert et al., 2010). All code is available at https://github.com/sjbeckett/DilutionMethod-ViralLysisEstimation and is archived at (Beckett and Weitz, 2018).

### 2.4 Nonlinear dynamics of virus-host interactions in the bottle

#### 2.4.1 Baseline model

The densities of phytoplankton, grazers and viruses are represented as *P, G* and *V* respectively. We can represent bulk population dynamics of these groups as:

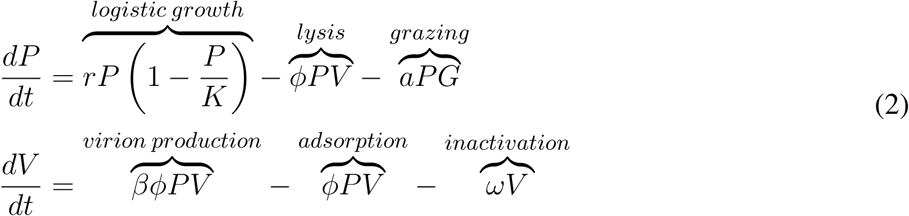

where we assume that the population of grazers stays constant within the time-scales of the dilution experiment (i.e. 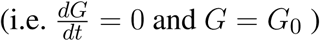 and *G* = G_0_). Here phytoplankton grow logistically with an intrinsic maximal growth rate *r* up to a carrying capacity *K*, are ingested by grazers at rate *a* and infected by viruses at rate *ϕ*. We assume, for now, that viral induced lysis is instantaneous creating a burst size, *β*, of viral progeny and that infective virions become inactive at rate *ω*. The baseline model, described by equation 2, has an initial viral-induced lysis rate given by:

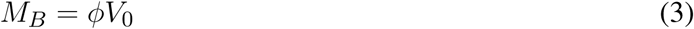

where *V*_0_ indicates the viral concentration at the time of sampling.

#### 2.4.2 Extension 1: considering infection dynamics

In the baseline model described in equation 2 viruses instantaneously lyse cells upon infection and there is no latent period. Here, we introduce an infected cell state, denoted I, which represents infected cells that can produce viral progeny. We assume that grazers are unable to differentiate between susceptible and infected cells, which are grazed upon at equal rates. The dynamics of this system are given as:

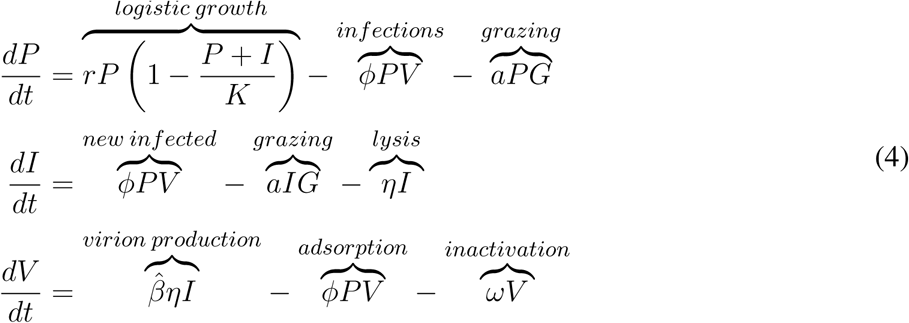

where we set

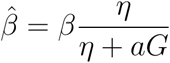

so that the average burst size in the explicit infection model is equal to that in the baseline model (Weitz, 2015). In the previous model phytoplankton cells were directly lysed. Here, it is only the phytoplankton cells in the infected class which lyse. As such, the rate of viral-induced lysis in the sample *M*_*I*_, that we wish to estimate is found as the rate of lysis from infected cells as a proportion of the total cell concentration (both infected and sucseptible cells):

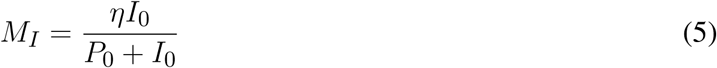

where *I*_0_ and *P*_0_ denote the sampled concentrations of infected and susceptible cells.

**Table 1.** Life history trait ranges used in the *in silico* dilution experiments used to create example and scanning plots.

| Symbol | Description | Value/Range | Units |
| --- | --- | --- | --- |
| $P$ | Density of susceptible phytoplankton cells | variable | cells ml<sup>-1</sup> |
| $I$ | Density of infected phytoplankton cells | variable | cells ml<sup>-1</sup> |
| $V$ | Density of infectious viral particles | variable | virions ml<sup>-1</sup> |
| $G_0$ | Density of grazers | 1000 | grazers ml<sup>-1</sup> |
| $r$ | Intrinsic per capita growth rate of phytoplankton | 1 | day<sup>-1</sup> |
| $K$ | Carrying capacity of phytoplankton | $2.2 \times 10^7$ (or scanned) | cells ml<sup>-1</sup> |
| $\phi$ | Adsorption rate of viruses to phytoplankton | $10^{-8}$ | ml/(virions·hour) |
| $\beta$ | Burst size | 50 | virions/cell |
| $\omega$ | Viral inactivation rate | 0.48 | day<sup>-1</sup> |
| $a$ | Grazer filtering rate | $2 \times 10^{-6}$ (or scanned) | ml/(grazers·hour) |

#### 2.4.3 Extension 2: effects of strain level diversity

We extend the baseline model presented in equation 2 to include viral and phytoplankton strain diversity as:

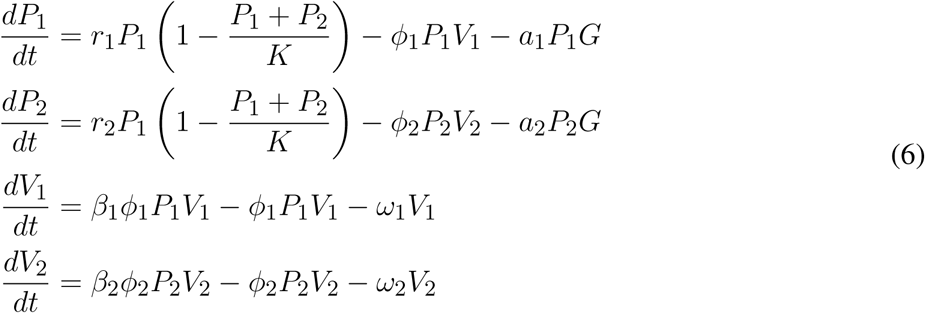

where both phytoplankton populations are limited by the same environmental conditions. For simplicity we assume that phytoplankton differ in growth rate, and that viruses only differ in their ability to adsorb to their respective hosts. Therefore we assume both viruses have the same burst size, β_1_ = β _2_ = β, and inactivation rates, ω _1_ = ω _2_ = ω. We also assume that grazer clearance rates are the same regardless of prey type (*a***_1_** = *a***_2_** = *a*). The range of life-history traits evaluated are shown in Table 2. We consider this model in two ways. First, we assume that the experimenter can tell the difference between the two phytoplankton types *P***_1_** and *P***_2_** and is able to count them separately. In this case the individual rates of viral-induced lysis, *M*_1_ and *M*_2_, are found respectively as:

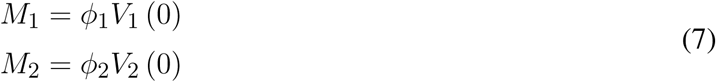

where *V*_1_. (0) and *V*_2_ (0) are the concentrations of each virus type at sampling time. On the other hand, if the experimenter is unable to identify differences between the phytoplankton types and counts them together, then the community wide bulk averaged rate of viral lysis, *M*_*c*_, with this model at sampling time is:

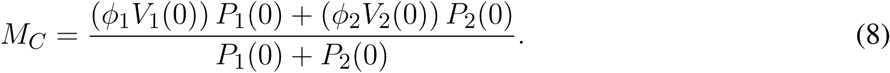

We choose to evaluate the perfomance of both community wide and individual viral lysis rate estimates. As before we assume that sampled concentrations are taken at the steady state for the system described in equation 6 given in Supplementary Information A.

**Table 2.** Life history trait values used in the *in silico* dilution experiments. Phytoplankton type 1 and virus type 1’s traits are varied within the given range, whilst phytoplankton type 2 and virus type 2’s traits are kept fixed.

| Symbol | Description | Value/Range | Units |
| --- | --- | --- | --- |
| $P_1$ | Density of phytoplankton type 1 cells | variable | cells ml<sup>-1</sup> |
| $P_2$ | Density of phytoplankton type 2 cells | variable | cells ml<sup>-1</sup> |
| $V_1$ | Density of infectious viral type 1 particles | variable | virions ml<sup>-1</sup> |
| $V_2$ | Density of infectious viral type 2 particles | variable | virions ml<sup>-1</sup> |
| $G_0$ | Density of grazers | 1000 | grazers ml<sup>-1</sup> |
| $r_1$ | Intrinsic per capita growth rate of $P_1$ | 0.1 – 10 | day<sup>-1</sup> |
| $r_2$ | Intrinsic per capita growth rate of $P_2$ | 0.2 | day<sup>-1</sup> |
| $K$ | Carrying capacity of phytoplankton | $2.2 \times 10^7$ | cells ml<sup>-1</sup> |
| $\phi_1$ | Adsorption rate of $V_1$ to $P_1$ | $10^{-10} - 10^{-7}$ | ml/(virions·hour) |
| $\phi_2$ | Adsorption rate of $V_2$ to $P_2$ | $10^{-10}$ | ml/(virions·hour) |
| $\beta$ | Burst size | 50 | virions/cell |
| $\omega$ | Viral inactivation rate | 0.48 | day<sup>-1</sup> |
| $a$ | Grazer filtering rate | $4.8 \times 10^{-5}$ | ml/(grazers·day) |

## 3 RESULTS

### 3.1 Analytical expressions for dilution growth rates

The difference in slopes between the classical and modified dilution curves is expected to be the viral lysis rate. We can check this expectation using the proposed dynamical model (equation 2). By calculating the instantaneous per capita phytoplankton growth rates for each incubation experiment at a particular dilution level *F* we find analytical forms for each of the respective dilution series:

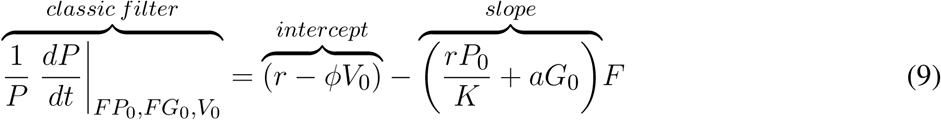

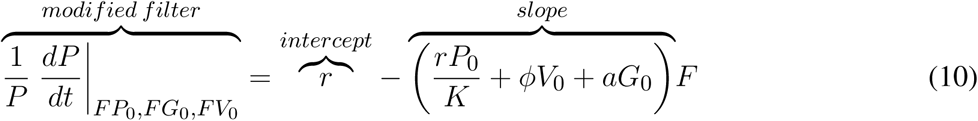

where *P*_0_, *G*_0_ and *V*_0_ are the initial sampled densities of phytoplankton, grazers and viruses. The viral induced lysis rate (ϕ *V*_0_) in the sample can be calculated as the difference between the slope values (or equivalently the intercepts) of the two dilution curves (Kimmance and Brussaard, 2010). We also consider an alternative approach - the VDiM - in which only viruses are diluted. Following the viral dilution approach, one can also estimate viral lysis as the slope from the corresponding dilution curve:

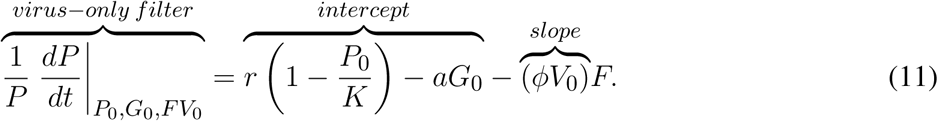

This suggests the viral dilution approach could be used as an alternative or complementary step in estimating viral lysis rates. Given these analytical expressions, we would expect the MDiM and the VDiM to perform well in the *instantaneous* limit. But inferences based on instantaneous growth rate measurements may not correspond well to growth rates based on measurements collected after a 24 hour incubation, during which the populations of viruses, in addition to cells, will deviate from their initial values.

We attempted to derive analytical expressions for each of the model extensions. For the model including an infected class (equation 4) our analytical expressions were unable to predict that we should recover viral lysis (see Supplemental Information D). This is due to the fact that the expression for total phytoplankton (susceptible and infected) per capita growth rates does not explicitly depend on the concentration of viruses, rather the rate of lysis is dependent on the concentration of infected cells. Whilst we were unable to recover the rate of viral-induced lysis using analytical expressions, we do not use this as evidence against the performance of dilution based approaches. Rather, this shows that even limited changes to nonlinear responses can make what may appear to be straightforward predictions difficult to analyse. In the second model extension, where diversity is examined, we were able to derive analytical expressions for the dilution curves. The case in which the experimenter can distinguish between phytoplankton types is trivially similar to that given for the baseline model. In the case in which the experimenter cannot distinguish between phytoplankton types we find theoretical expectations for the classic dilution series curve as:

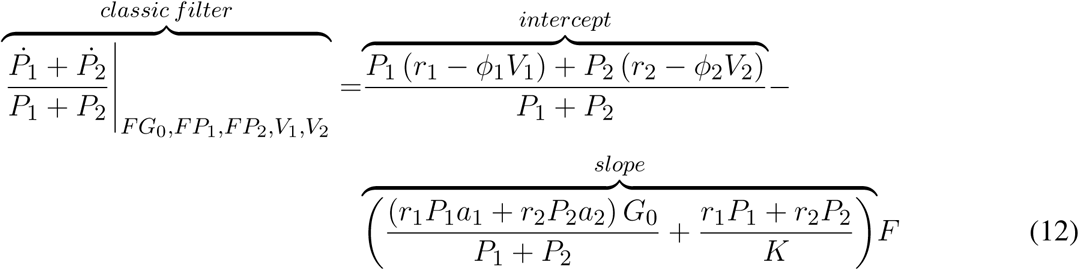

and for the modified dilution series:

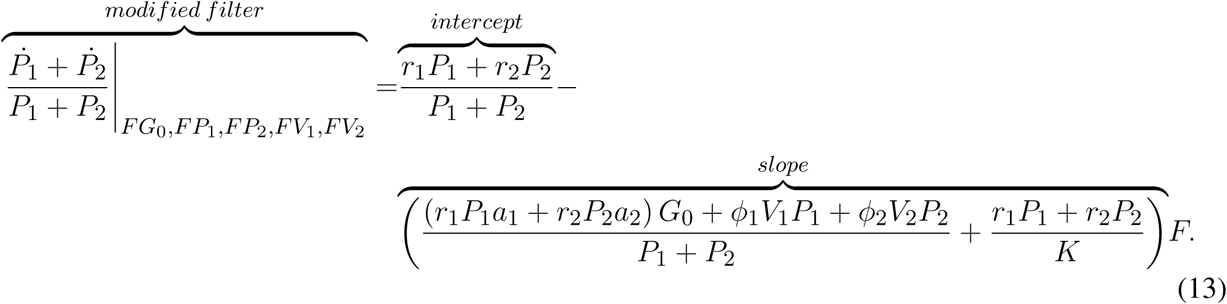

The difference in both the intercepts and the slopes of these series (equations 12 and 13) is equal to the bulk-average rate of viral lysis in the community:

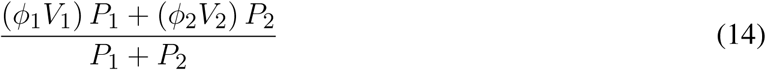

as previously defined in equation 8. Similarly for the VDiM, we find:

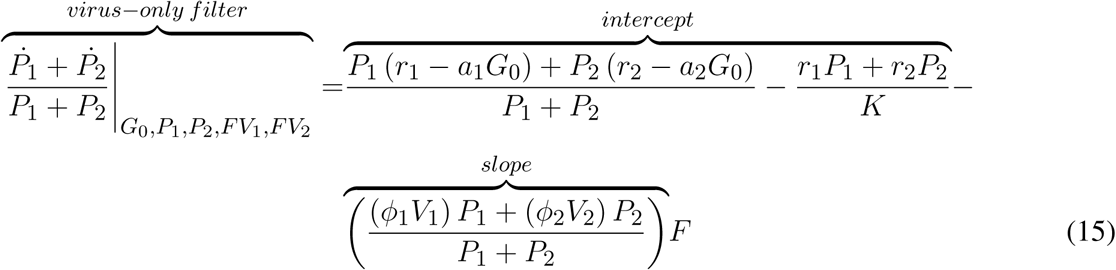

where the slope is predicted to estimate the bulk-average rate of viral lysis for the community, as defined in equation 8. As such, the expectation is that we should be able to infer rates of viral lysis at both the type- and community-level for the system described in equation 6.

Whilst these analytical expressions match our expectations under instantaneous measurement, the ability to which they are able to do so may differ depending on the life-history traits exhibited in the microbial community. Whilst growth increases the phytoplankton population, this can be limited through niche competition and by the top-down controls of grazing and viral lysis. At steady state these limitation processes are equal to growth (Figure 2). The ability to estimate viral lysis rates might depend on which of these limitation mechanisms is dominant. We now use *in silico* dilution experiments to test and evaluate these expectations.

**Figure 2.**
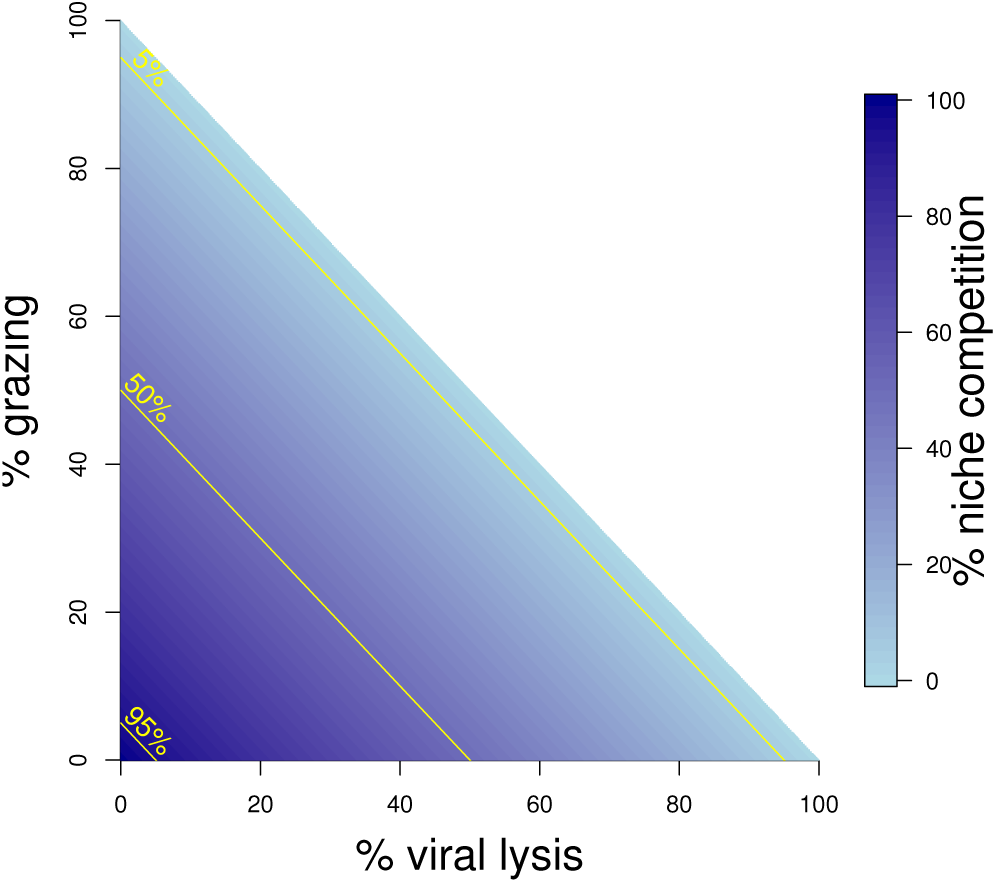
Schematic representing mortality processes affecting the phytoplankton. The sum of three mortality processes: grazing, viral-induced lysis and niche competition must add to 100%. Grazing and lysis are indicated on the axes, whilst niche competition is indicated by shading. 5%, 50% and 95% isoclines of niche competition are labelled.

### 3.2 Evidence of potential bias in lysis rate estimates

Figure 3 shows the rate estimates for growth, grazing and viral lysis made by the CDiM, MDiM and VDiM following a 24h incubation of the population dynamics described in equation 2. Figure 3a shows the population dynamics of phytoplankton cells and viruses from time 0h to 24h (as indicated by arrows) within individual incubation bottles in each of the three types of dilution series. This phase portrait shows that the dynamical trajectories of viruses and cells differ across incubation bottles, but also that they oscillate around the systems fixed point - in our simulations the sampled density of viruses and cells. Apparent growth rates were calculated for each incubation bottle in each of the classic, modified and viral dilution series using equation 1 and are plotted in Figure 3b with the best fitting linear regression. All three dilution curves appear to be linear. Using the intercept and slope values calculated from the dilution curves as defined by the CDiM, MDiM and VDiM protocols we compare inferred ecological rates to model input rates in Figure 3c. Note that only the MDiM provides estimates of all rates simultaneously; and that the CDiM and MDiM estimate grazing the same way and are therefore equal. For this set of parameters the CDiM appears to underestimate growth rates, whilst grazing rates appear to be overestimated. Both the MDiM and the VDiM infer high rates of viral lysis. However, the MDiM underestimates the viral lysis rate. The mechanistic basis for these biases in estimates of lysis rates can be understood in terms of the nonlinear dynamics that arise in 24 hrs (see Figure 3a) vs. those expected given the theory of instantaneous lysis. The signal of these nonlinear dynamics is not apparent from analysis of the dilution curves.

**Figure 3.**
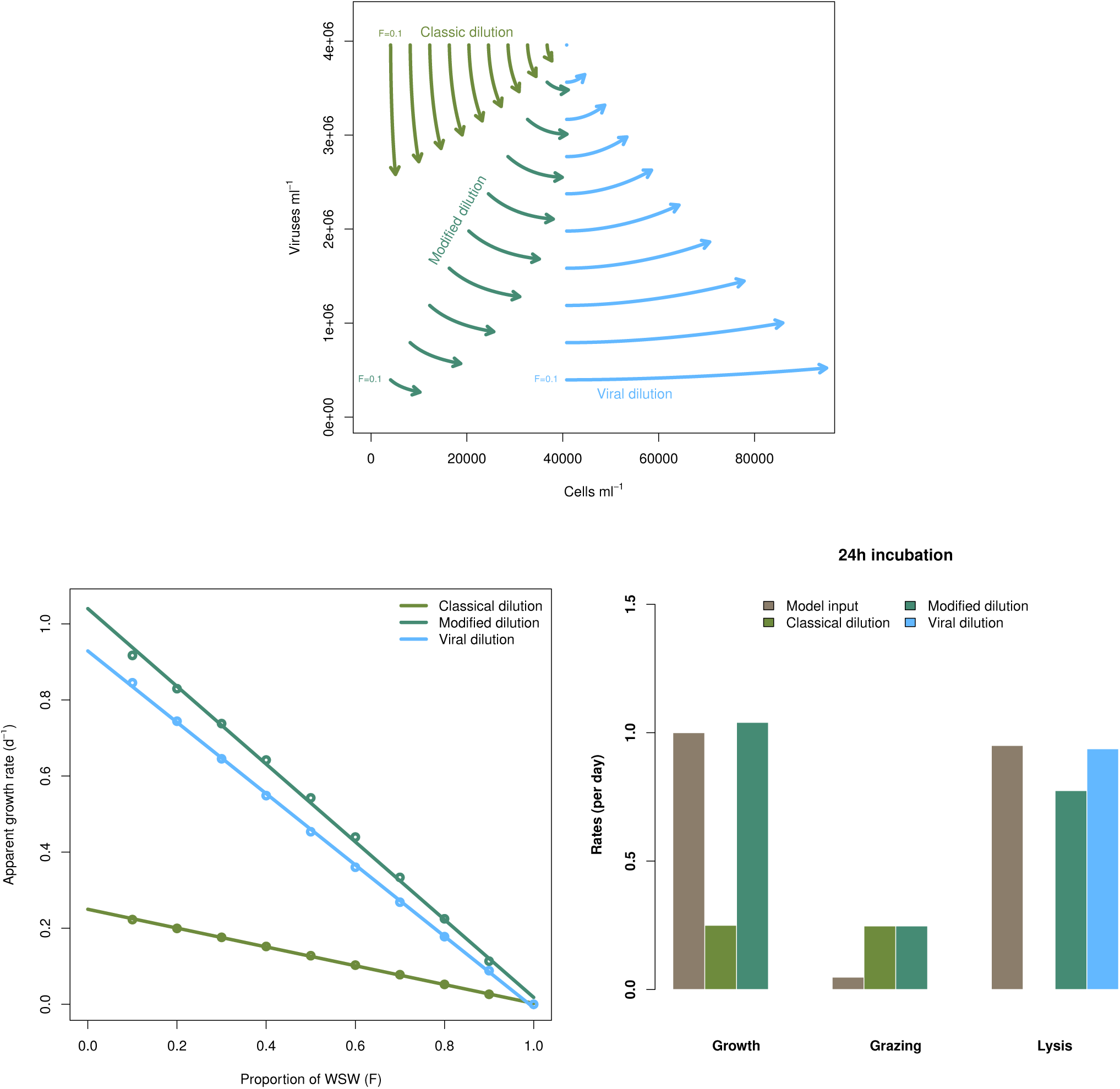
Inference of ecological rates using dilution methods. (a) Population dynamics of phytoplankton cells and viruses within individual incubation bottles from the classical, modified and virus dilution series between 0h and 24h. The intital sampled concentrations of cells and viruses (*F* = 1) are shown as the single point. (b) Dilution curves constructed from calculations of apparent growth rates within each of the incubation bottles at 24h. (c) Comparing model rate inputs to dilution-based rate estimates of growth, grazing and viral induced lysis derived from the dilution curves.

### 3.3 Robustness of lysis rate estimates to variation in life history traits

To explore the robustness of viral-induced lysis rate estimation we examined different parameter regimes using two approaches. In the first, we changed the relative amounts of bottom-up to top-down mortality, as well as the ratio between viral-induced and grazing mortality (consistent with the indicated isoclines of niche competition in Figure 2) which are shown using 24h incubations in Figure 4. This indicates that both the MDiM and the VDiM performed better when bottom-up mortality i.e. niche competition is low. The VDiM performed best when niche competition was low, but the MDiM provided better estimation when niche competition was high. The relative amount of top-down mortality partitioned between viral lysis and grazer did not appear to change the estimation bias associated with the different methods.

**Figure 4.**
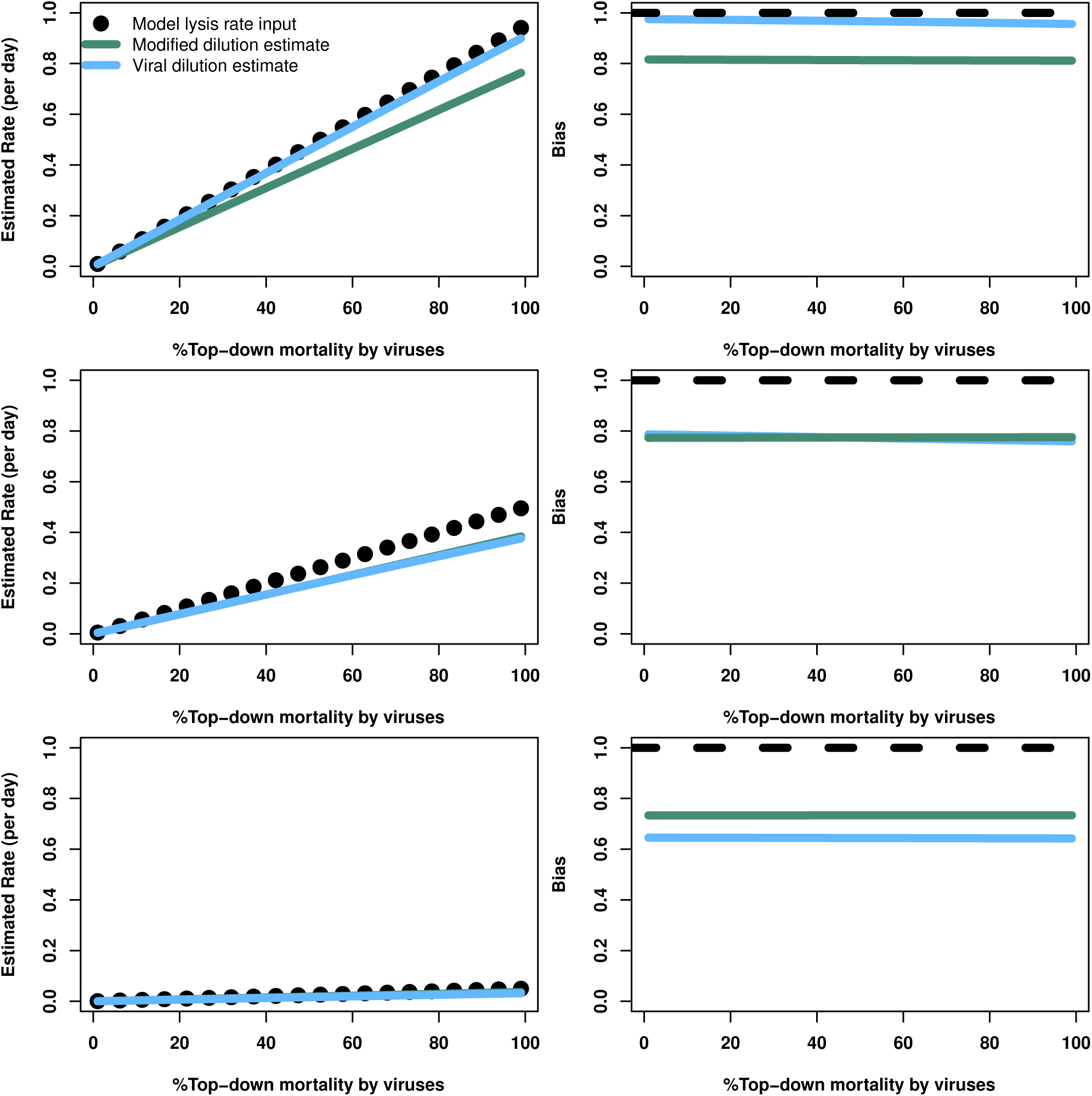
Rate estimates (left) and bias (right) of viral lysis rate in the baseline model following 24h incubation. Each row shows a different level of niche competition as indicated by the lines across Figure 2 (top: 5%, middle:50%, bottom: 95% niche competition).

The analytical results suggest that both the modified and viral dilution methods should work well under near-instantaneous measurement. Hence, we repeated this procedure using a shorter incubation period to see its effect on viral lysis estimation. Figure 5 shows that using a 2h incubation period dramatically improves estimation ability across all conditions. Again, we see that the VDiM appears to provide better estimates than the MDiM when niche competition is low, but the reverse is true when niche competition is high.

**Figure 5.**
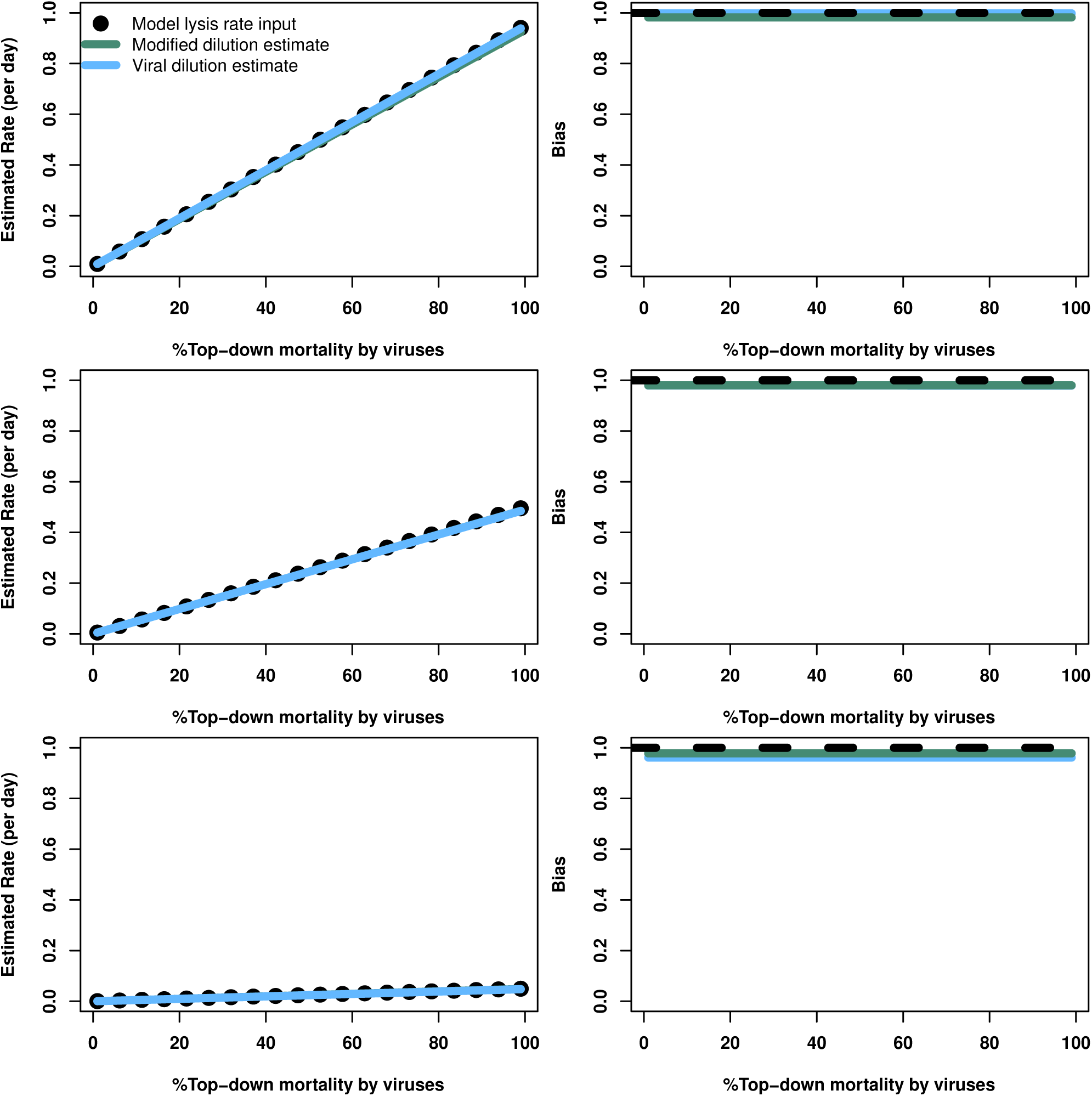
Rate estimates (left) and bias (right) of viral lysis rate in the baseline model following 2h incubation. Each row shows a different level of niche competition as indicated by the lines across Figure 2 (top: 5%, middle:50%, bottom: 95% niche competition).

In order to further address the robustness of the two inference methods to differences in life-history traits and model parameterisation we used a Latin Hypercube sampling design to assess estimation ability from an ensemble of model simulations which were assessed using short (2h) and long (24h) incubations. The parameter ranges are shown in Table 3. These results suggest shorter incubations may improve estimation of viral-induced lysis and that the VDiM is potentially more robust across systems with different life-history traits (Figure 6). However, we note that there was a large variation in the efficacy of both methods and that the VDiM could erroneously report a negative lysis rate estimate 6b.

**Table 3.**
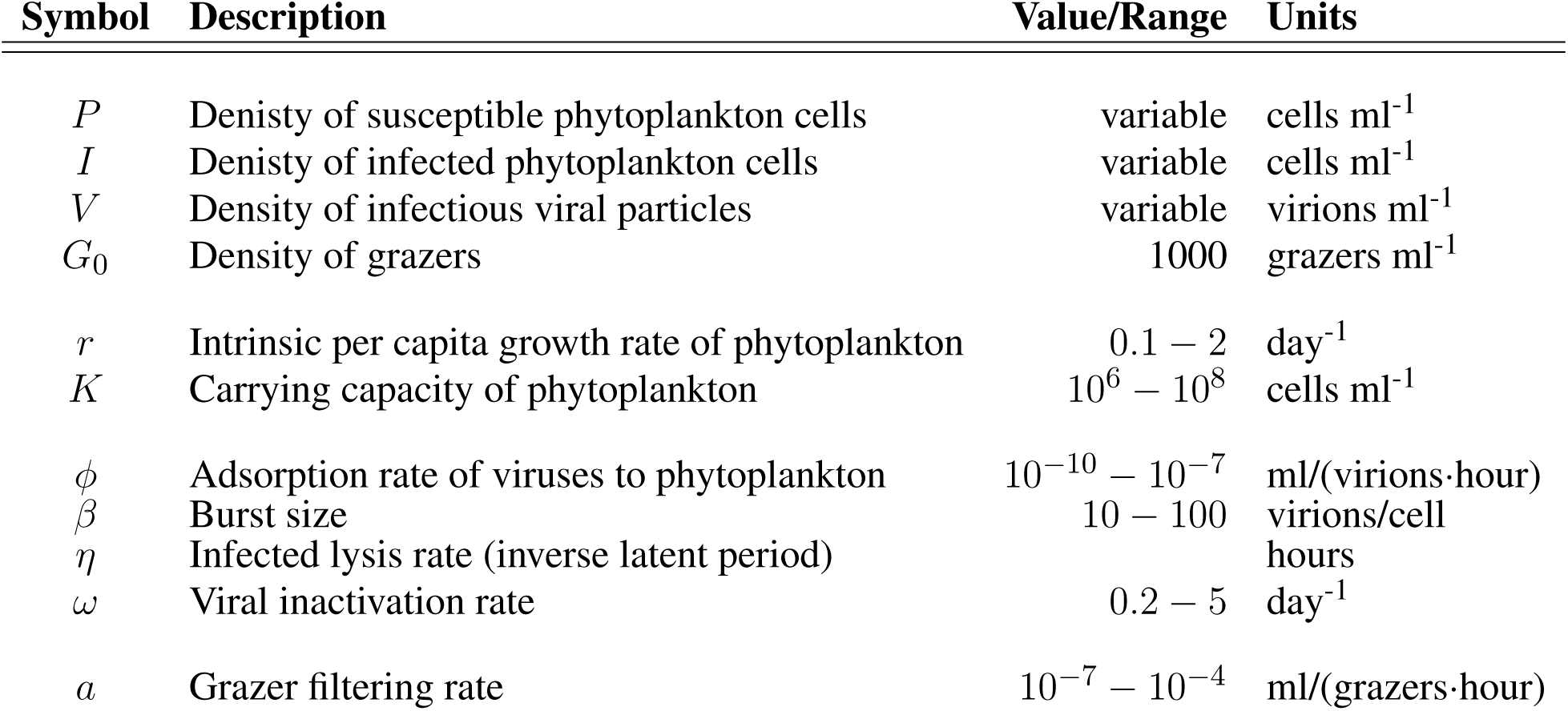
Life history trait ranges used in the *in silico* dilution experiments used to create density plots. Parameter choices were made from these ranges using a random latin hypercube sampling design.

**Figure 6.**
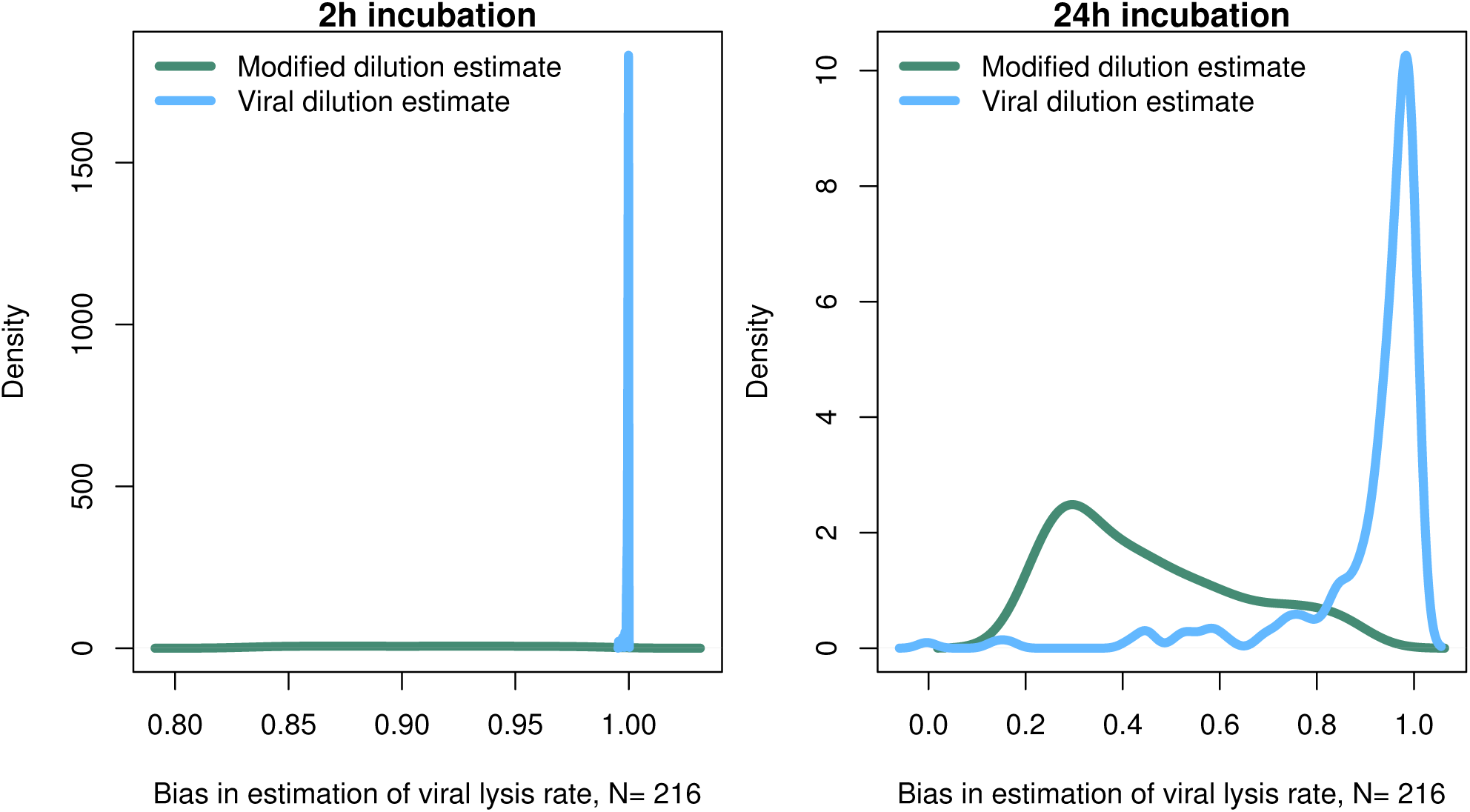
Density of estimation bias in the baseline model made by the MDiM and VDiM for incubations of 2h (left) and 24h (right). Density plots are from N simulations (out of 500) from random latin hypercube sampled parameter choices.

### 3.4 Robustness of lysis rate estimates given variation in viral latent periods

The results from the baseline model (Figures 3 - 6) suggest that viral-induced lysis rate estimation may be improved by using shorter incubations and, in some instances, the viral dilution method. However, the baseline model assumes that viral adsorption leads to instantaneous cellular lysis and release of new infectious virions. Using the extended model that includes infected phytoplankton (equation 4) we ask whether the latent period duration affects inference of viral lysis rates. We chose to examine this model using three different latent periods: 15 minutes, 4 hours and 24 hours.

The results from a system with a short 15 min latent period (Figure 7, S1 and S4) are qualitatively similar to the baseline model. However, the efficacy of both the MDiM and the VDiM is reduced relative to that found under the assumptions of the baseline model - this is particularly apparent in comparing the estimation bias under a short incubation (Figure S1 relative to Figure 5). This can be seen firstly by the drop in observed variation of estimation bias following a 2h incubation for a 4h latent period (Figures 8 and S2) and a 24h latent period (Figures 9a and S3). Additionally, as can be seen in Figure 8a and Figure 9 when the latent period exceeds the incubation period, the estimates from both the MDiM and the VDiM are not only low, but also quantitatively similar. During 24h incubations the VDiM appears more robust than the MDiM, Figures 8b-9b, but both methods had large variation in estimation bias.

**Figure 7.**
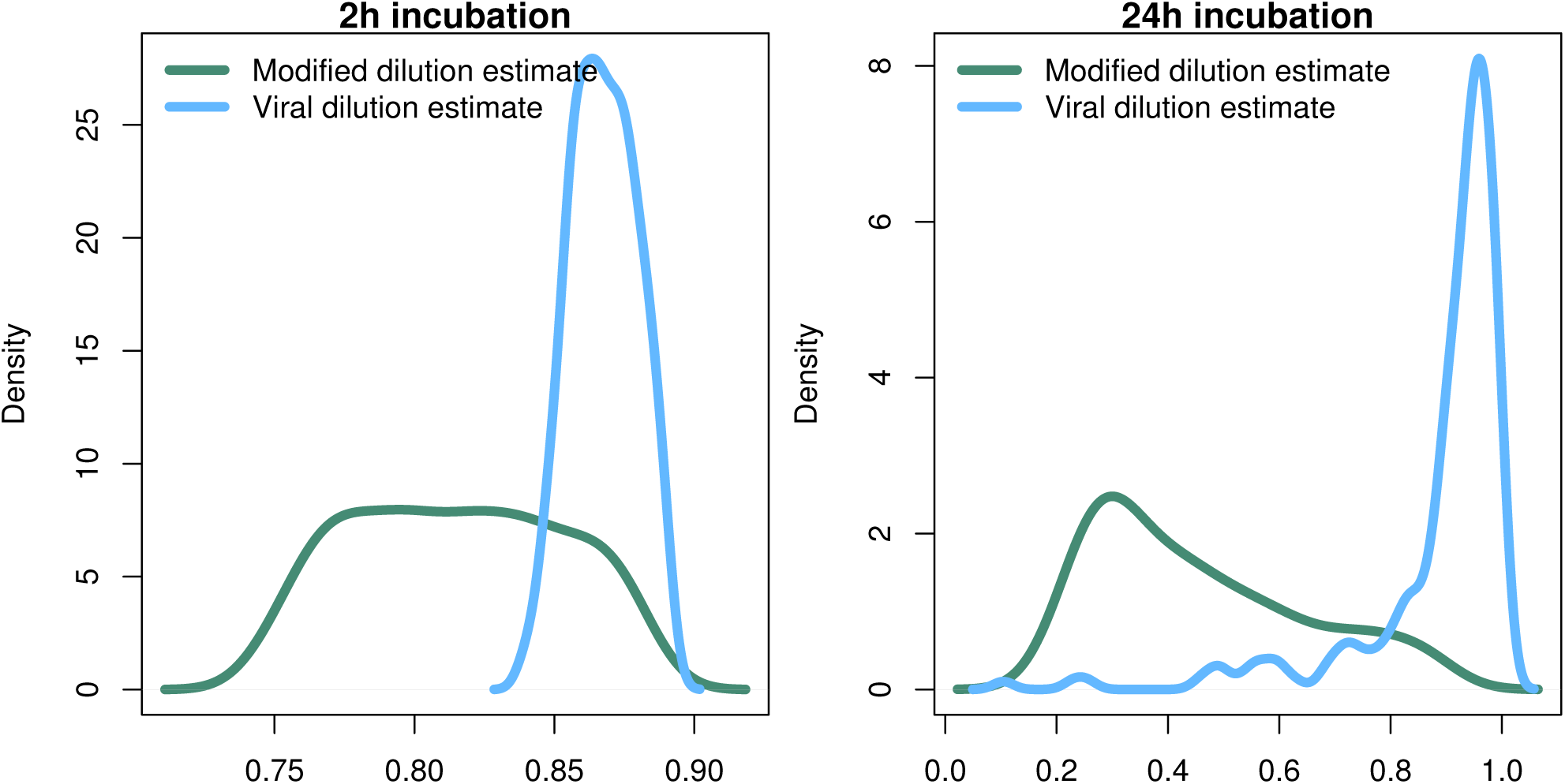
Density of estimation bias in the infected class model, with 15 min latent period, made by the MDiM and VDiM for incubations of 2h (left) and 24h (right). Density plots are from N simulations (out of 500) from random latin hypercube sampled parameter choices.

**Figure 8.**
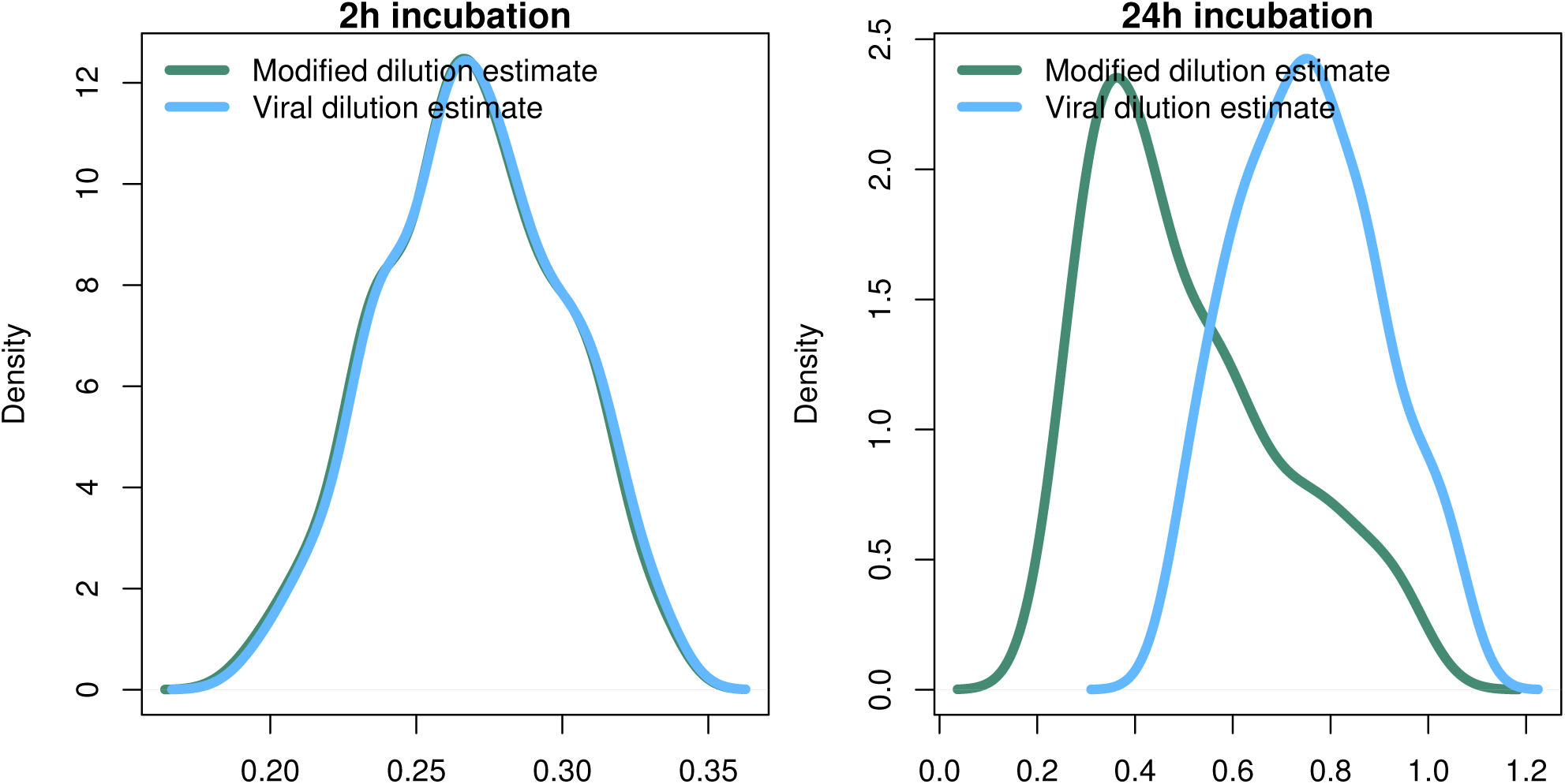
Density of estimation bias in the infected class model, with a 4 hour latent period, made by the MDiM and VDiM for incubations of 2h (left) and 24h (right). Density plots are from N simulations (out of 500) from random latin hypercube sampled parameter choices.

**Figure 9.**
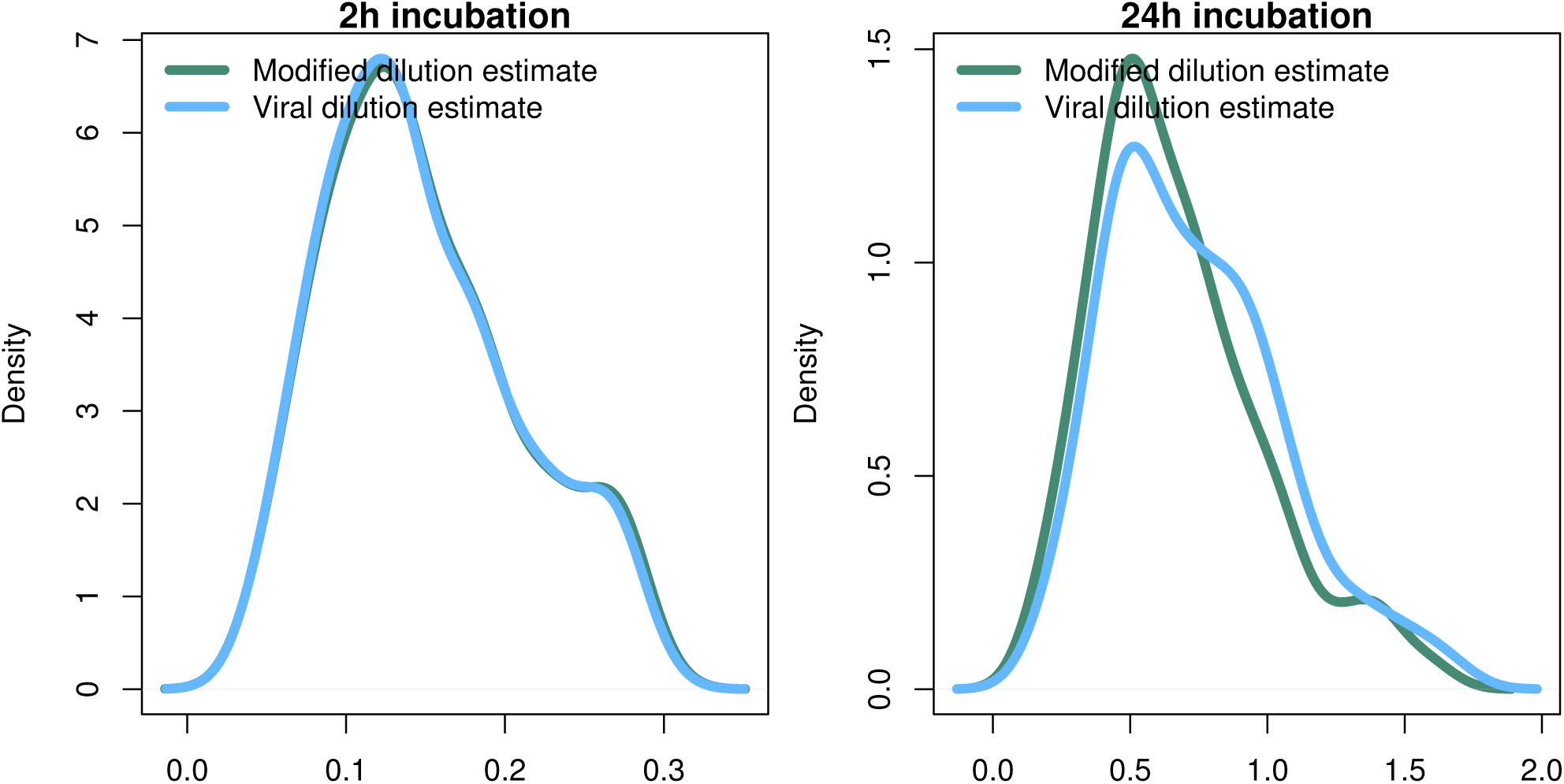
Density of estimation bias in the infected class model, with a 24 hour latent period, made by the the MDiM and VDiM for incubations of 2h (left) and 24h (right). Density plots are from N simulations (out of 500) from random latin hypercube sampled parameter choices.

### 3.5 Strain level diversity may effect the ability to estimate viral-induced lysis

Life-history trait differences between the members of interacting microbial communities could lead to biased measurements within dilution method experiments. To highlight this potential effect we consider a community with two phytoplankton types P1 and P2, which are each infected by a strain-specific virus, V1 and V2. We further assume that P1 has a faster growth rate relative to P2, but has a lower ambient steady-state concentration, as shown in Figure 10. This is an expectation of Kill-the-Winner dynamics in which faster growing phytoplankton are capable of supporting a larger virus population via negative density-dependent selection (Thingstad, 2000; Zhao et al., 2013). Following the dilution of populations (time 0 h) type P1 is able to recover much faster than type P2. If the observer is unable to distinguish between these two phytoplankton types a different dynamic is apparent and may lead to misleading interpretations of viral-lysis in the community. In Figure 10b the model input rates of viral-lysis and the corresponding estimates of viral-induced lysis are shown at the level of the community and at type-level. Here, viral-induced lysis was underestimated at the type levels for both phytoplankton. However, the community viral-induced lysis rate was overestimated.

**Figure 10.**
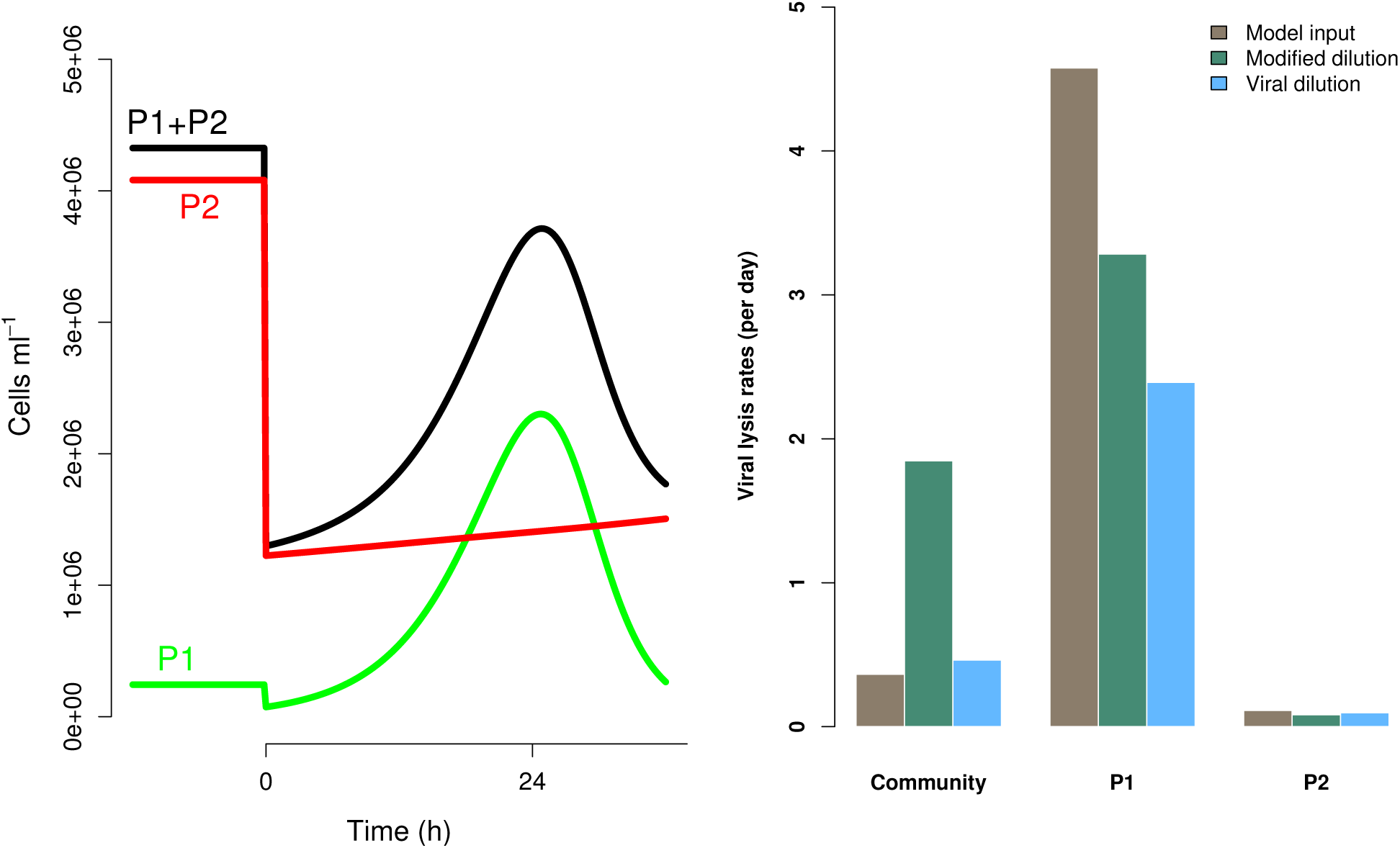
Differences between strains may hinder efforts to estimate lysis rates. (a) Phytoplankton population dynamics during an incubation experiment in the modified dilution series with *F* = 0.3. Here phytoplankton type P1 grows much faster than type P2, but has a lower steady state abundance (population level before t=0h). (b) The viral lysis rate model input and estimated rates for the system shown in (a). Viral lysis rates are measured at the community- and type-level.

To explore the ability of the modified and viral dilution methods to robustly estimate viral-induced lysis rates in phytoplankton communities we fixed the life-history traits of phytoplankton type P2 whilst varying the ambient concentration and growth rates of phytoplankton P1. The results are shown following a 2h incubation in Figure 11 and following a 24 h incubation in Figure 12. Similarly to the baseline model we find that estimation ability improves with shorter incubation periods, but we note that this model does not include an infected class and lysis is instantaneous. Community estimates of viral-induced lysis may be overestimated or underestimated depending on the life-history traits in the community. These effects are larger during 24h incubations than 2h incubations. Following a 2h incubation the 95% quantiles of estimation bias reported in Figure 11 and 12 are (0.98,1.34) for the MDiM and (1.00,1.21) for VDiM; but following a 24h incubation are (−0.14, 5.63) for MDiM and (0.14,1.69) for VDiM. The regimes in which overestimation and underestimation occur are also dependent on observation timing. At the type-level (bottom two rows) we see both the MDiM and the VDiM generally underestimate viral-induced lysis rate across the variation in life-history traits.

**Figure 11.**
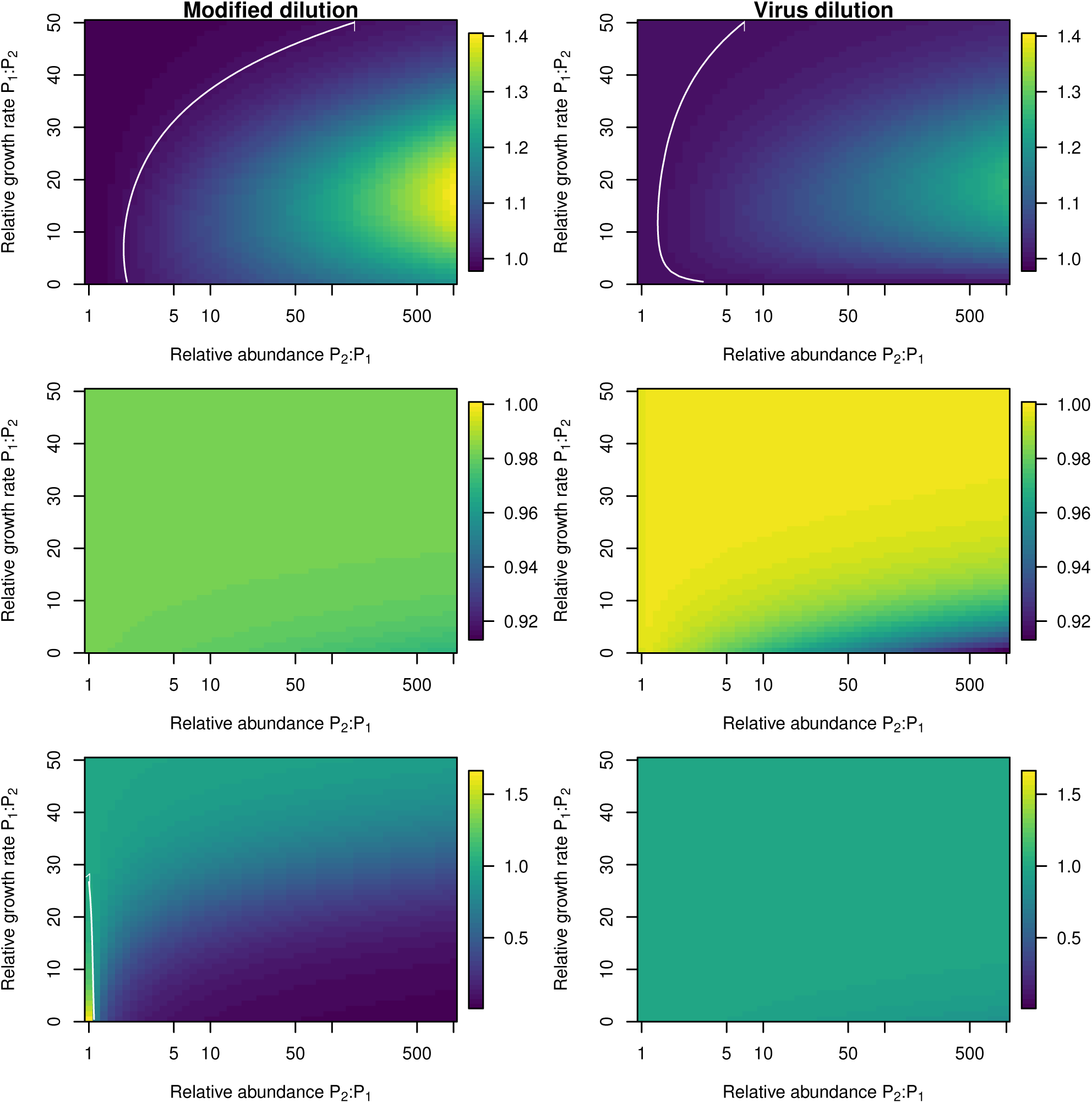
Estimation bias in viral lysis rates by the modified dilution method (left) and viral dilution method (right) following a 2h incubation. Rows show community-wide estimates (top), *P*_1_ estimates (middle) and *P*_2_ estimates (bottom). White contours indicate an estimation bias equal to one, where the estimated rate is equal to the model input.

**Figure 12.**
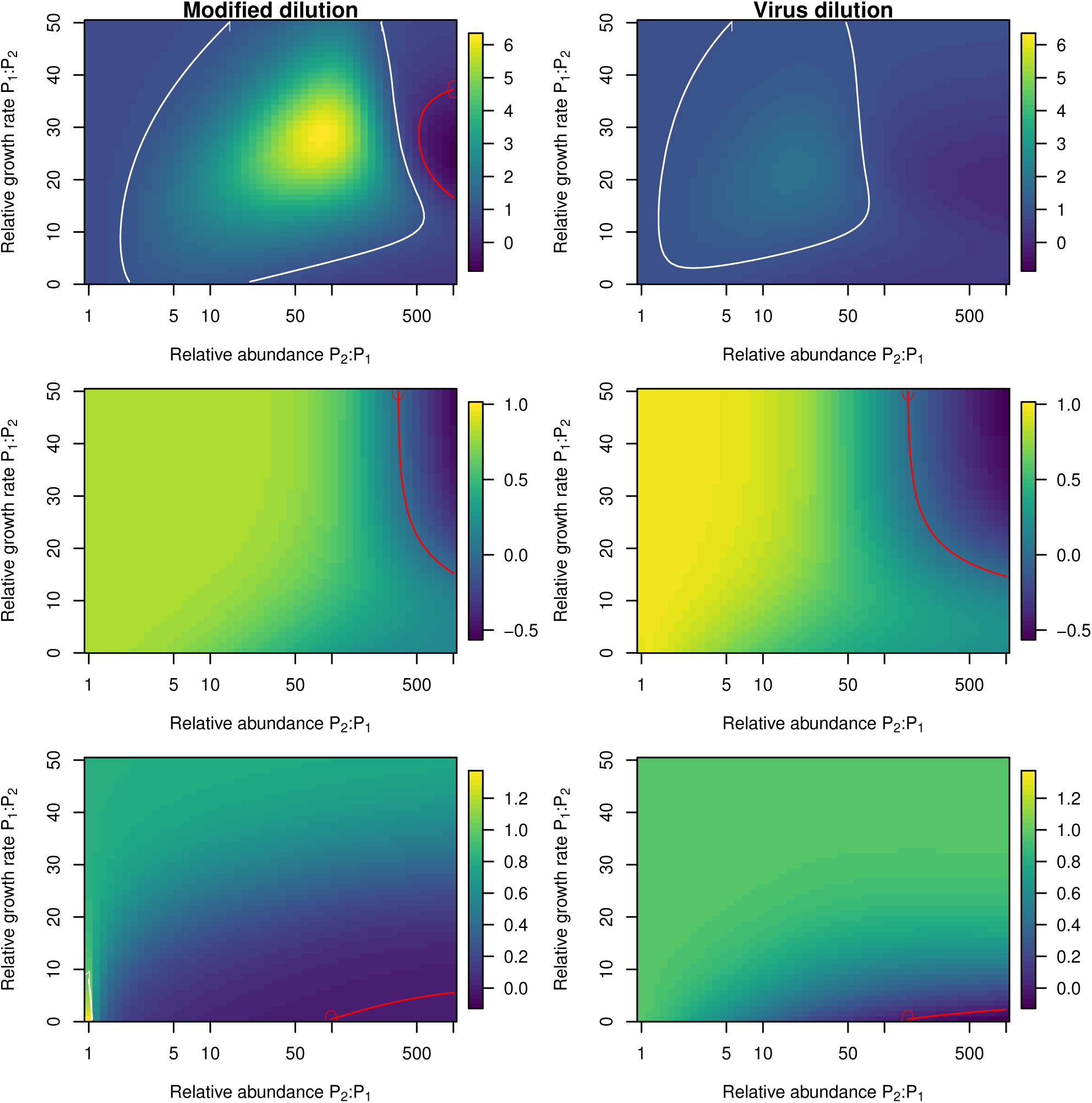
Estimation bias in viral lysis rates by the modified dilution method (left) and viral dilution method (right) following a 24h incubation. Rows show community-wide estimates (top), *P*_1_ estimates (middle) and *P*_2_ estimates (bottom). White contours indicate an estimation bias equal to one, where the estimated rate is equal to the model input. Red contours indicate an estimation bias of zero. This shows where dilution slopes go from negative to positive such that estimates of viral lysis rates take negative values.

## 4 DISCUSSION

We have systematically analyzed the potential for dilution-based methods to infer viral lysis rates of phytoplankton. In doing so we have combined nonlinear models of community dynamics in a specific experimental context. We derived equations for expected dilution curves (equations 9 and 10) which provides a principled basis for why the difference in these slopes may be able to approximate viral lysis. We were also able to recover the rate of viral lysis when analysing the case of viral dilution (equation 11), under the knowledge that this analysis represents instantaneous expectations, and not those after a long incubation period. However, our in silico simulations suggest viral lysis rate may be difficult to measure in practice.

Our simulation results suggest that estimation ability of viral-lysis rates is poorer under increased bottom-up control via niche competition. This complements previous research in which we found that niche competition might also reduce the ability to estimate grazing rates (Beckett and Weitz, 2017). This suggests greater understanding of the nutrient and incubation conditions and how they relate to the physiology of plankton communities in dilution experiments is necessary. Nutrient addition is commonly used to alleviate nutrient limitation within dilution experiments. As highlighted by Calbet and Saiz (2017) different nutrient addition treatments are applied within different dilution experiments. As well as asking how nutrient addition effects niche competition, future consideration should be given to how particular organisms will differentially respond to a particular nutrient addition strategy (Lagus et al., 2004) and how this affects the ability to learn about *in situ* ecological processes.

We investigated two potential approaches to improve estimates of viral-induced lysis rates - reducing incubation length in the MDiM and applying a new, VDiM in which only virus concentrations are diluted. Reducing incubation timing appeared to improve estimation ability under circumstances when the incubation length was greater than the infection latent period. This suggests *a priori* knowledge of latent periods could help improve experimental design - however such *a priori* knowledge may not be available *in situ*. Our simulations do not model how ecological processes may be effected by diel forcing e.g. (Arias et al., 2017), which could be an additional complicating factor in attempting to use dilution based approaches to estimate ecological rates of phytoplankton, grazers and viruses.

The VDiM did appear to have improved efficacy relative to the MDiM when populations were not limited by bottom-up control. In order to practically implement the VDiM we see two possibilities. First, empiricists could attempt to resuspend viruses e.g. using flocculation techniques (John et al., 2011; Poulos et al., 2018). A different approach to achieving a gradient of viral dilution could be to use the filtrate (as opposed to the diluent) of the classical dilution filtered water and mixing this with whole seawater at different proportions.

Figure 3 represents a cautionary tale for the field in the interpretation of dilution experiments. Observations made using dilution experiments typically find similar rates of phytoplankton growth and grazing mortality e.g. (Morison and Menden-Deuer, 2017). However, as shown in Figure 3 it is important to remember the ecological context of such measurements - growth rate estimates made by the classical dilution method are limited by the activity of viruses and hence may be underestimated. We caution empiricists to consider the role of viruses in future experiments.

In addition to treating phytoplankton and viruses as bulk entities, we investigated how well dilution based estimates perform when diversity is included at both type- and community-level. We found that fast growing strains have the potential to recover quickly and therefore dominate the apparent lysis rate, even if they are relatively small contributors to the true signal. This level of virus-host interaction complexity remains over-simplified. Virus-host interactions are expected to be highly specific (though some viruses have been observed to infect across phyla (Malki et al., 2015)). Nonetheless, at the strain level there may exist a range of specialist to generalist virus types (Weitz et al., 2013). In addition we continue to treat grazers as a bulk entity, though Calbet and Saiz (2013) find that trophic chains can affect the results of dilution experiments. Grazing is also assumed to be non-preferential which is an assumption that could be challenged (Wirtz, 2014; Pasulka et al., 2015). The structure of the network of interactions between viruses, phytoplankton cells and grazers will affect both the observed population dynamics and the expected individual type- and community-level rates of mortality. Investigating how well simple approximations, such as those made by bulk-population models of the dilution method, work in more complex ecological communities warrants further investigation.

There are a number of additional assumptions that could prove limiting to the dilution method. Inherent in our assumptions are that life-history traits are constant and do not vary in time or with changing environmental conditions during the incubation time. Additionally we assume that grazing and viral infection processes are linearly affected by dilution (i.e. Holling type I) which may be a simplistic assumption. Saturation in grazing responses (Li et al., 2017), or viral infection (Kimmance and Brussaard, 2010) could lead to nonlinear dilution curves which need different methods of interpretation. Without observations of the population dynamics between the beginning and end of the dilution experiments it may be difficult to assess the appropriateness of the conceptual mechanistic framework from which dilution-based rate estimates are inferred. We assume that the filters used by the classical, modified and viral dilution series all perform perfectly which may not be the case e.g. Pasulka et al. (2015), and that nutrient levels are not reduced by dilution (though see (Pasulka et al., 2015)). Additional challenges arise from consideration of nutrient regeneration via viral lysis, lysogeny, the potential for preferential grazing on infected cells, removal of free viruses by grazing (Kimmance and Brussaard, 2010) and mixotrophy (Caron, 2016). These could all serve as routes for future study.

We have shown that even with perfect measurement ability, large uncertainty exists in the ability of dilution based methods to estimate rates of viral-induced lysis. These uncertainties are related to mismatch between the expectation of exponential recovery from dilution for the bulk community and predicted nonlinear dynamics and time-delayed feedbacks within complex microbial communities of viruses, grazers and their microbial prey. Improving estimates of viral effects *in situ* requires that we revisit the mechanistic assumptions of experimental protocols in light of the increasing understanding of the diversity of marine microbial and viral communities.

## AUTHOR CONTRIBUTIONS

SJB and JSW designed the research and revised the manuscript. SJB performed the simulations, analysis and wrote the manuscript.

## ACKNOWLEDGMENTS

We thank Will Ratcliff, David Caron, Debbie Lindell, Joey Leung and Mick Follows for feedback that has improved this manuscript. We thank Daniel Muratore for conducting a review of the simulation code.

